# Co-aggregation and parallel aggregation of specific proteins in major mental illness

**DOI:** 10.1101/2023.06.12.544321

**Authors:** Bobana Samardžija, Maja Juković, Beti Zaharija, Éva Renner, Miklós Palkovits, Nicholas J. Bradshaw

**Author notes:** BS and MJ are joint first authors. Corresponding author: Address: Sveučilište u Rijeci, Odjel za biotehnologiju, Radmile Matejčić 2, 51000 Rijeka, Croatia.

## Abstract

**Background:** Disrupted proteostasis is an emerging area of research into major depressive disorder. Several proteins have been implicated as specifically forming aggregates in the brains of subsets of patients with psychiatric illnesses, these proteins include CRMP1, DISC1, NPAS3 and TRIOBP-1. It is unclear, however, whether these normally aggregate together in the same individuals, and, if so, whether each protein aggregates independently of each other (“parallel aggregation”) or if the proteins physically interact and aggregate together (“co-aggregation”).

**Materials and methods:** Post mortem insular cortex samples from major depressive disorder and Alzheimer’s disease patients, suicide victims and control individuals had their insoluble fractions isolated and tested by Western blotting to determine which of these proteins are insoluble, and therefore likely aggregating. The ability of the proteins to co-aggregate (directly interact and form common aggregate structures) was tested by systematic pairwise expression in SH-SY5Y neuroblastoma cells, examined by immunofluorescent microscopy.

**Results:** Many individuals displayed multiple insoluble proteins in the brain, although not enough to imply interaction between the proteins. Cell culture analysis revealed that only a few of the proteins analyzed can consistently co-aggregate with each other: DISC1 with CRMP1 and TRIOBP-1. DISC1 was able to induce aggregation of full length TRIOBP-1, but not its domains expressed individually.

**Conclusions:** While specific proteins are capable of co-aggregating, and appear to do so in the brains of individuals with mental illness, and potentially also with suicidal tendency, it is more common for such proteins to aggregate in a parallel manner, through independent mechanisms.

## Introduction

Major depressive disorder (MDD), bipolar disorder and schizophrenia are all severe and often chronic mental illness, which have profound influences on patients, their families and society in general. The underlying pathophysiology of these conditions is partially understood, in large part due to advances in uncovering genetic risk factors, however these studies present a highly heterogenous picture, with many risk factors of small effect accounting for a proportion of heritability, with remaining risk assumed to come from rarer mutations (1-3). As a supplement to this approach, we and others have proposed instead to consider the role of proteinopathy in chronic mental illnesses (4). Specifically, in partial analogy to how specific proteins form misfolded or unfolded aggregates in the brains of patients with neurodegenerative disease, similar aggregates, of differing proteins, may also exist in the brains of at least some patients with chronic mental illness. Unlike in neurodegenerative diseases, we do not expect these aggregates to be neurotoxic.

Studies investigating aggregation of these proteins in mental illness generally use insolubility as an indicator of aggregation (4). Protein aggregates are larger agglomerates with either an incorrect or random structure, that are normally insoluble in the cell and common experimental systems. By taking homogenized brain samples, purifying out only the more insoluble protein fraction and testing by Western blotting, it can be determined whether a specific protein is insoluble in the original brain sample (5, 6). Should a normally soluble protein be found in the insoluble fractions of patient brain samples specifically, then this provides a strong argument for this protein aggregating in the associated illness. Such approaches have detected Disrupted in Schizophrenia 1 (DISC1), dysbindin 1, Collapsin Response Mediator Protein 1 (CRMP1) and TRIO Binding Protein isoform 1 (TRIOBP-1) as insoluble in a subset of patients with schizophrenia, bipolar disorder and/or MDD (5-10), while Neuronal PAS protein 3 (NPAS3) is implicated through an aggregation-inducing mutation (11, 12). In all cases, insoluble protein has also been detected in mammalian cell culture models and found to be equivalent to visible protein aggregates in the cell body (6, 7, 9, 10, 12, 13). The fact that these events are not diagnosis specific, however, raises the interesting possibility that protein aggregation, both generally or of these specific proteins, may be a common feature in mental illness. To date, however, sufficient samples sizes have not been available to determine whether these aggregation events correspond to specific subtypes or symptoms of these illnesses.

Another interesting finding from previous work is that in some, but not all, cases, insoluble DISC1 co-exists in the same brain samples as insoluble dysbindin-1 and CRMP1 (7, 9). Subsequent analysis then showed that DISC1 could directly bind to and induce aggregation of both of these proteins, in a process of “co-aggregation”. Other combinations of proteins implicated in major mental illness have not yet been tested in the same way, leaving it unclear whether, in general, these proteins each aggregate alone, possibly in distinct patient populations, or co-aggregate together as part of a more general proteinopathy. In favor of this latter idea is the fact that the ubiquitination and proteasome system of the cell has been shown to be generally less functional in schizophrenia patients, while total insoluble protein is higher (14-16). It is therefore also possible that common causes in the cellular environment could lead to multiple proteins aggregation in the same brain, or even the same neuron, without physically interacting with each other, in a process of “parallel aggregation”.

We therefore investigate here the potential for co-aggregation and parallel aggregation in major mental illness, using two distinct approaches: investigation of protein insolubility in a set of human brain samples, and systematic investigation of co-aggregate forming potential in human neuroblastoma cells.

## Materials and Methods

### Human brain tissue

Collection, storage and distribution of human brain tissue was approved by the Committee of Science and Research Ethics of the Ministry of Health of Hungary (No. 6008/8/2002/ETT) and the Semmelweis University Regional Committee of Science and Research Ethic (No. 32/1992/TUKEB). Use of brain tissue for this project was approved by the Ethical Committee of the University of Rijeka, Department of Biotechnology (23-10-2018). All work was performed in accordance with the Declaration of Helsinki and all national and European laws. Informed familial consent or legal permission was acquired before collecting each sample. As part of the Hungarian Lenhossék program, brains were collected with short post-mortem delay (2 -10 h), and samples of different brain regions were isolated using the “micro-punch” technique (17, 18). During each step of the microdissection procedure, brains were frozen and then stored at -80 °C. For this work, such samples were used from the insular cortices of victims of suicide (n = 15) and control individuals (n = 11), as well as patients with MDD (n = 4) and Alzheimer’s disease (n = 3). Levels of NPAS3 in this cohort have been investigated previously, and demographic information can be found in that publication (13). All tissue donors were Hungarian, of whom 44% were female and 56% male.

### Insolubility assay

Samples of brain homogenate had their insoluble protein fraction purified as described previously (6). Briefly, samples of 10% brain homogenate are solubilized, treated with DNaseI and then subjected to a number of ultracentrifugation steps, after which the soluble (liquid) protein fraction is discarded. The insoluble pellet is then resuspended and centrifuged again. Buffers variously include high salt, high sucrose and detergents, to isolate only the most insoluble protein in the sample, which is predicted to contain protein aggregates. Cell lysates had their insoluble proteins isolated using a similar technique, also described previously (13).

### Antibodies

Primary antibodies were purchased against β-actin (Origene, Rockville, MD, USA, OG-TA811000), CRMP1 (ProSci, Poway, CA, USA, 3625), DISC1 (Thermo Fisher Scientific, Weltham, MA, USA, 40-6800), Flag (Merck, Darmstadt, Germany, F1804), GFP (Merck, G6795) and TRIOBP-1 (Atlas Antibodies, Stockholm, Sweden, HPA019769). Secondary antibodies were purchased from Thermo Fisher Scientific (31430, 65-6120 and A11037).

### Plasmids

Vectors encoding human CRMP1 (9) and TRIOBP-1, both full length and fragments (19), were gifts from Prof. Dr. Carsten Korth (Heinrich Heine University, Düsseldorf, Germany), with CRMP1 then being subcloned into pENTR1A no ccDB (20) (Dr. Eric Campeau, AddGene clone 17398, Watertown, MA, USA). Vectors encoding NPAS3 (13) and a TRIOBP-1 aggregation-resistant mutant (6) were described previously. Gateway entry vectors encoding full length human DISC1 and NPAS3, came from the ORFeome Collaboration (22, 23) (DNASU Plasmid Repository, clones HsCD00516321 & HsCD00080332, Tempe, AZ, USA). Entry vectors were transferred into pDEST-CMV-N-EGFP (24) (Prof. Robin Ketteler, AddGene clone 122842) and/or pdcDNA-FlagMyc (B. Janssens, BCCM/LMBP Plasmid Collection, clone LMBP 4705, Zwijnaarde, Belgium) using LR Clonase II (Thermo Fisher Scientific). More details of the plasmids and primers used are in supplementary tables S1 and S2. All plasmids were confirmed by sequencing.

### Cell culture

HEK293 human kidney cells were cultured in D-MEM (Thermo Fisher Scientific), supplemented with 5% HyClone Cosmic Calf serum (Cytiva, Marlborough, MA, USA) penicillin and streptomycin (Pan-Biotech, Aidenbach, Germany). Plasmids were transfected into cultured cells using Metafectene (Biontex, Munich, Germany) according to manufacturer’s protocols. SH-SY5Y human neuroblastoma cells were cultured in D-MEM/F-12, supplemented with 5% fetal calf serum (both Thermo Fisher Scientific), non-essential amino acids, penicillin and streptomycin (Pan-Biotech). Plasmids were transfected into cultured cells using Metafectene Pro (Biontex) according to manufacturer’s protocols.

### Western blotting

Samples were denatured in 156 mM Tris pH 6.8 / 5% SDS / 20 mM DTT / 25% glycerol with bromophenol blue for 5 minutes at 95 °C and then separated on bis-acrylamide gels. Gels were transferred to PVDF membranes (Macherey-Nagel, Düren, Germany) using a Transblot Turbo system (Bio-Rad, Hercules, CA, USA) and transfer was confirmed by staining with 0.5% Ponceau S / 2% acetic acid. Membranes were blocked for 1 hour at room temperature in PBS / 0.05% Tween-20 / 5% milk powder, and then stained overnight using primary antibodies diluted in the same buffer. Membranes were washed 4 times over 30 minutes with PBS / 0.05% Tween-20 and then stained with secondary antibodies in the same buffer. Membranes were then washed 4 times over 30 minutes with PBS / 0.05% Tween-20, and the signal revealed using ECL (Thermo Fisher Scientific). The signal was detected and quantified using a ChemiDoc MP Imaging System and associated software (Bio-Rad).

### Immunocytochemistry and microscopy

Cells on glass coverslips were fixed with PBS / 4% paraformaldehyde for 15 minutes, and then permeabilized with PBS / 0.5% Triton X-100 for 10 minutes. Cells were blocked with PBS / 10% goat serum (Merck) for 30 minutes and then stained with primary antibody in the same media for 2-4 hours. Cells were then washed 3 times (5 minutes each) with PBS, and then stained with secondary antibodies and DAPI (Merck) in PBS / 10% goat serum for 1 hour. Cells were washed three more times and affixed to slides with Fluoroshield histology mounting medium (Merck). Cell were viewed on am IX83 inverted microscope (Olympus, Shinjuku, Japan) and images taken using an Orca R2 digital CCD camera (Hamamatsu Photonics, Hamamatsu, Japan) and cellSens software (Olympus). For quantified analysis of protein (co-)aggregation, tubes containing plasmids were coded and randomized. The researcher who transfected these into cells, and then analyzed the proportion of cells with (co-)aggregation, was therefore blinded as to which plasmid(s) each cell was expressing. Only after quantification were the samples decoded for statistical analysis using GraphPad Prism.

## Results

### Presence of multiple insoluble proteins in individual brain samples

In order to investigate possible co-aggregation or parallel aggregation of proteins implicated in mental illness, a cohort of insular cortex samples were collected, consisting of victims of suicide (n = 15), control individuals (n = 11) and smaller numbers of patients with MDD (n = 4) or Alzheimer’s disease (n = 3). The insular cortex was used because of its previous association with a variety of neurological and psychiatric disorders (26).

These samples were homogenized and the insoluble protein fractions of each sample was then purified, and these fractions were investigated by Western blotting, to determine if insoluble (aggregating), CRMP1, DISC1 and/or TRIOBP-1 was present in them (figures 1A-D and S1). The original, non-purified, brain homogenates were also Western blotted for comparison (figure S2). Some level of insoluble protein was seen in many of the samples, with a few instances of individuals showing very high amounts of one specific insoluble protein. Notably one depression patient showed very high levels of insoluble DISC1 aggregation (figure 1A and S1B) and one suicide victim similarly showed high levels of insoluble TRIOBP-1 (figure 1B and S1A, major 72 kDa species), while individuals of various diagnoses expressed high levels of CRMP1 (both the long variant, Lv, and short variant, Sv) as insoluble proteins (figure 1C and S1C, Lv: 70 kda, Sv: 65 kDa). NPAS3 has previously been analyzed in these samples (13), with some individuals showing high levels of insoluble NPAS3 (figure 1D, major 120 kDa species).

**Figure 1.**
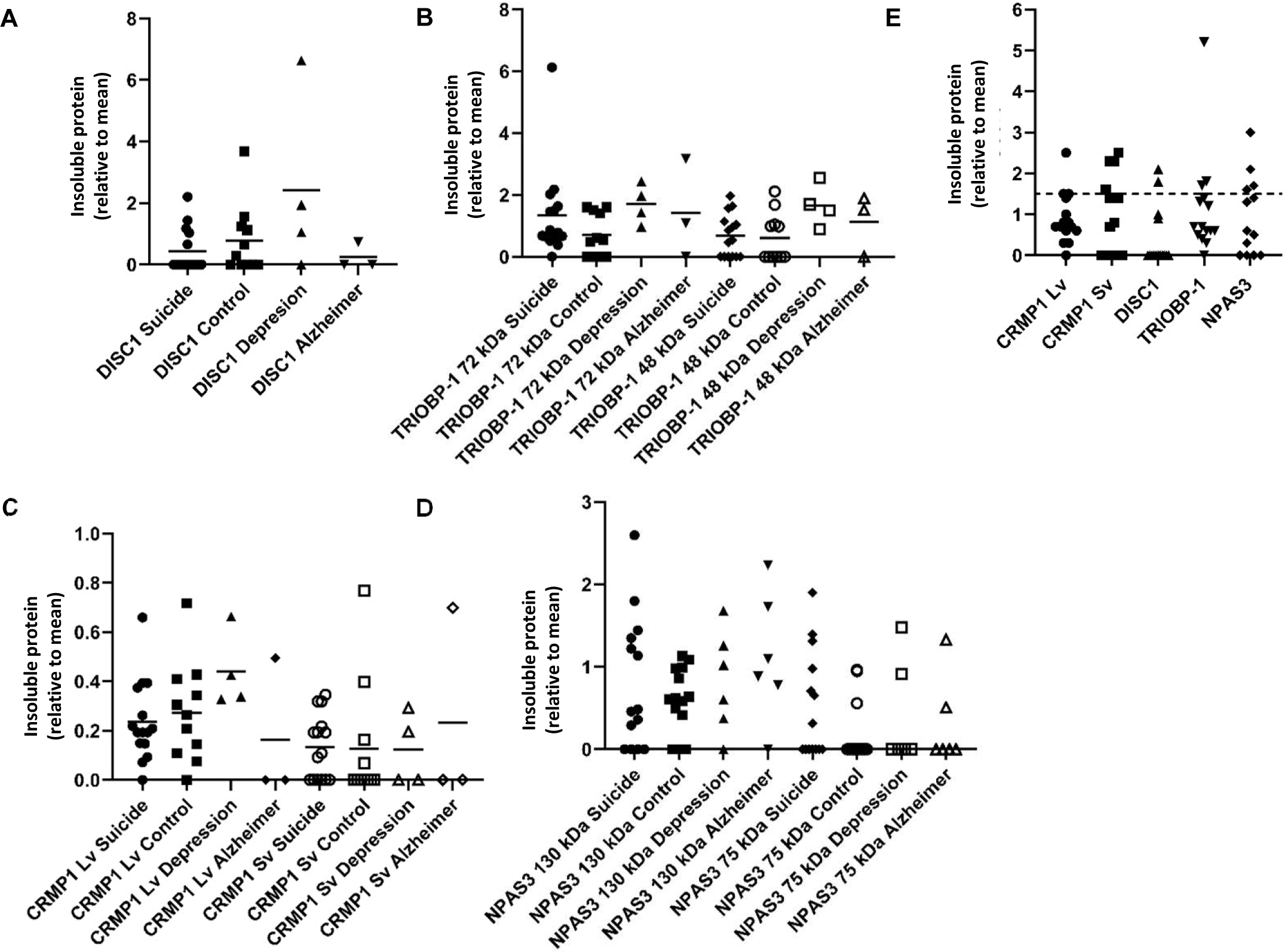
Insoluble protein in human insular cortex samples. **(A-D)** Quantified levels of DISC1 (A), TRIOBP-1 (B), CRMP1 (C) and NPAS3 (D) protein seen in the insoluble protein fraction of post mortem insular cortex samples. Values are normalized to a common sample loaded on each membrane. Original Western blot data for DISC1, TRIOBP-1 and CRMP1 are in figure S1. NPAS3 data (D) has been published previously (13), but is reanalyzed and summarized here for comparison. **(E)** For the protein isoforms analyzed further (CRMP1 Lv and Sv, DISC1 70 kDa, TRIOBP-1 72 kDa, NPAS3 130 kDa) the proteins present in the insoluble pellet at a level at least 50% higher than the mean, which are interpreted as aggregating for the purposes of this study. **(F)** The number of proteins found to be insoluble (aggregating) in each sample. Graphs prepared using GraphPad Prism.

In the majority of cases, levels of insoluble protein were not normally distributed, with many individuals showing little or none of a specific protein, and others showing higher levels. For the purposes of this experiment, we considered any sample that contained more than 1.5× the mean level of an individual protein to potentially contain that protein in an aggregating state (figure 1E). 9 of the 15 suicide victims contained at least one potential aggregating protein by this definition (60%), compared to 5 out of 11 control individuals (45%, figure 1E). The 4 MDD and 3 Alzheimer’s samples all contained potential aggregating proteins by this definition. Of these, more than half showed at least two proteins to be potentially aggregating, with some showing three and one MDD patient having four (figure 2). It therefore seems likely that multiple proteins do aggregate in subsets of patients. The proportion of individuals expressing multiple aggregating proteins suggests that these are more likely to be a result of each protein aggregating individually (parallel aggregation), rather than through the active effect of one aggregating protein on the aggregation state of another (co-aggregation). The proteins that were most commonly found to be insoluble together in samples were CRMP1 Lv, CRMP1 Sv and DISC1, however incidences of insoluble NPAS3 and TRIOBP-1 being present were also seen (figure 2). There was no obvious correlation between levels of insolubility of any two individual proteins (figure S3).

**Figure 2.**
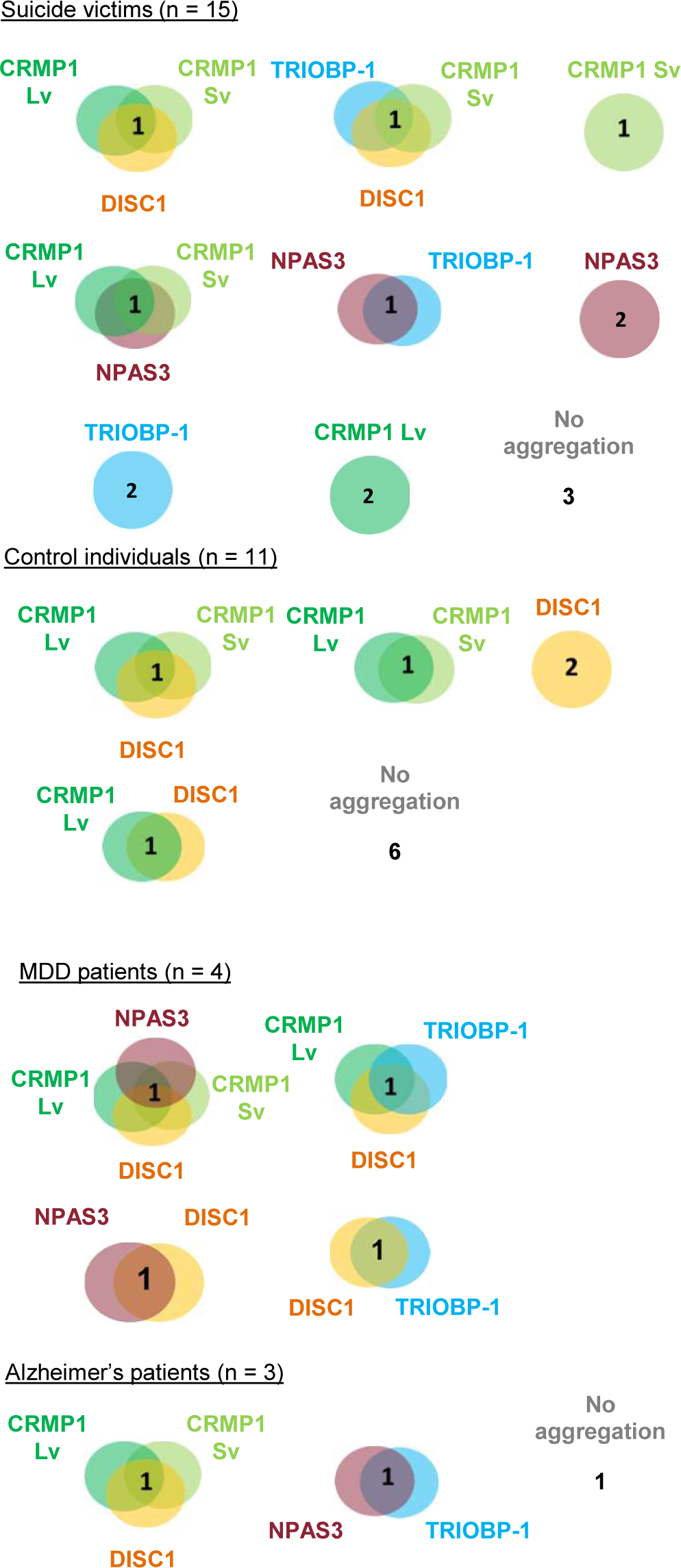
Detailed breakdown of co-occurrence of insoluble proteins in the insular cortex samples. Each circle or Venn diagram represents one or more individuals (as indicated by the number) and the proteins that are insoluble in the insular cortex sample from that individual. Individuals not shown did not show any insoluble proteins. Insoluble is defined as a level of a specific protein in the insoluble fraction that is at least 1.5x the average across all 37 samples tested. DISC1, NPAS3 and TRIOBP-1 represent the major 75 kDa, 130 kDa and 70 kDa species respectively.

### Pairwise co-expression studies in neuroblastoma cells show only DISC1 readily forms co-aggregates, with both CRMP1 and TRIOBP-1

Genes coding for CRMP1 (Lv and Sv), DISC1 (L isoform), NPAS3 and TRIOBP-1 were each expressed in two plasmid vectors, one that added a Flag tag to them, and one that fused them to EGFP. Protein expression was confirmed by Western blotting (figure 3A,B). These were then expressed in SH-SY5Y neuroblastoma cells, to establish whether or not they spontaneously formed visible aggregates. Using Flag-tagged proteins, CRMP1 was found in the cell body (Lv and Sv, figure 3C,D) and NPAS3 was primarily found in the nucleus (figure 3E, with occasional cells showing cytoplasmic localization instead). In contrast, DISC1 and TRIOBP-1 were each consistently seen to aggregate in the cell body (figure 3F,G). These results are all broadly as would be predicted from previous work (7, 9, 10, 13, 19). There were no changes in expression pattern of any of the Flag-tagged proteins when they were co-expressed with EGFP (figure S4A-F). EGFP-fused versions of these proteins behaved identically to their Flag-tagged counterparts (figure S4G-M) with the exception of EGFP-CRMP1, which was sometimes seen to form aggregate-like structures (a minority of cells expressing EGFP-CRMP1 Sv, a majority expressing EGFP-CRMP1 Lv, figure S4H-I).

**Figure 3.**
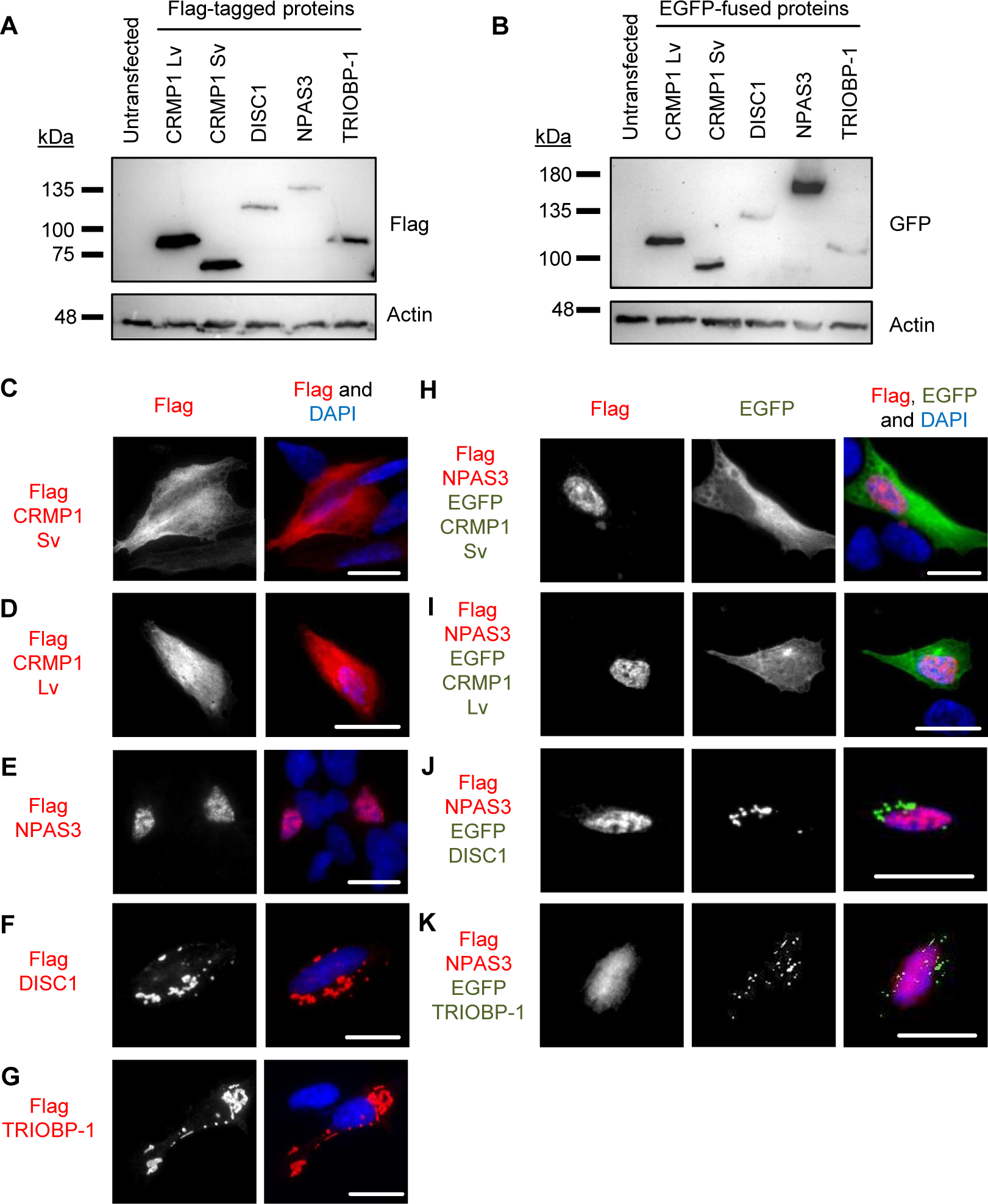
Systematic pairwise testing of co-aggregation in cell culture. **(A)** Western blots of Flag-tagged proteins used in these experiments, expressed in HEK293 cells. **(B)** Equivalent blot of EGFP-fused proteins used in this experiment, plus EGFP alone as a control. Remaining images show constructs expressed in SH-SY5Y neuroblastoma cells: **(C)** Flag-tagged CRMP1 Sv, not aggregating. **(D)** Flag-tagged CRMP1 Lv, not aggregating. **(E)** Flag-tagged NPAS3, not aggregating. **(F)** Flag-tagged DISC1, aggregating. **(G)** Flag-tagged TRIOBP-1, aggregating. **(H)** Flag-tagged NPAS3 and EGFP-fused CRMP1 Sv, neither aggregating. **(I)** Flag-tagged NPAS3 and EGFP-fused CRMP1 Lv, neither aggregating. **(J)** Flag-tagged NPAS3 and EGFP-fused DISC1, only DISC1 is aggregating. **(K)** Flag-tagged NPAS3 and EGFP-fused TRIOBP-1, only TRIOBP-1 is aggregating. All cell photos are typical of 3 or more independent experiments. Scale bars represent 10 μm. Figure S5 shows versions of the experiments in (H) to (K) using the reciprocal vectors.

Pairs of proteins were then systematically co-expressed in SH-SY5Y to determine whether they could co-aggregate in these cells. In all cases, experiments were performed with one protein being Flag tagged and the other fused to EGFP, and then a reciprocal experiment performed to verify the results with the other combination of vectors. Unless otherwise noted, all results were consistent, regardless of which protein had the Flag tag, and which was fused to EGFP. NPAS3 showed no consistent signs of co-aggregation with any of the other proteins (figures 3H-K, S5). Similarly, while TRIOBP-1 formed aggregates in cells containing CRMP1 Sv, there was no sign of co-aggregation (figures 4A, S6A). Co-aggregation of TRIOBP-1 and CRMP1 Lv was seen in some cells, but only when TRIOBP-1 was Flag-tagged and CRMP1 fused to EGFP, not in the reverse situation (figures 4B, S6B). In contrast, DISC1 was seen to co-aggregate with both variants of CRMP1, in many but not all cells with Sv, and in most cells with Lv (figures 4C-E, S6C-E). Subpopulations of CRMP1 Lv and Sv also co-aggregated with each other in a majority of cells examined (figure 4F, S6F). Notably, however, while both DISC1 and TRIOBP-1 were seen to each aggregate in almost all cells in which they were co-expressed, in some cells they were seen to clearly co-aggregate with each other, while in others they aggregated in parallel (figures 4G-H, S6G).

**Figure 4.**
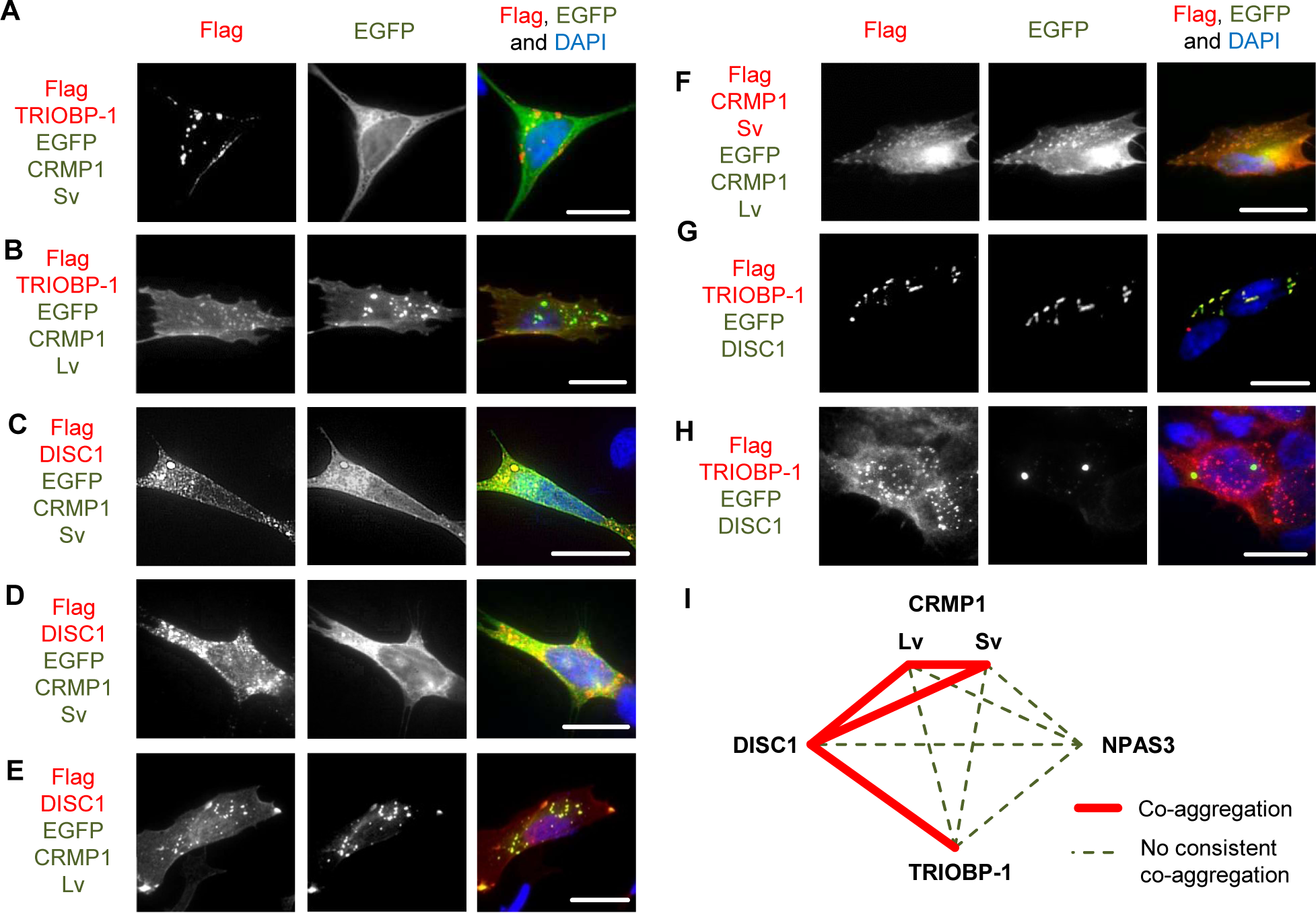
Further systematic pairwise testing of co-aggregation in cell culture. Images show constructs co-expressed in SH-SY5Y neuroblastoma cells: **(A)** Flag-tagged TRIOBP-1 and EGFP-fused CRMP-1 Sv, only TRIOBP-1 aggregates. **(B)** Flag-tagged TRIOBP-1 and EGFP-fused CRMP-1 Lv, only TRIOBP-1 aggregates. **(C-D)** Flag-tagged DISC1 and EGFP-fused CRMP-1 Sv, example of cell with co-aggregation (C) and DISC1 aggregation only (D). **(E)** Flag-tagged DISC1 and EGFP-fused CRMP-1 Lv showing co-aggregation. **(E)** Flag-tagged CRMP1 Sv and EGFP-fused CRMP1 Lv showing co-aggregation. **(G-H)** Flag-tagged TRIOBP-1 and EGFP-fused DISC1, example of cell with co-aggregation (F) and parallel aggregation (G). **(I)** Summary of co-aggregation results. All cell photos are typical of 3 or more independent experiments. Scale bars represent 10 μm. Figure S6 show versions of these experiments using the reciprocal vectors.

These results confirm one previous report of co-aggregation from the literature, that of DISC1 with CRMP1 Sv (7), and indicates two novel ones: that of DISC1 with TRIOBP-1 and CRMP1 Lv (figure 4H). All other pairs of proteins examined were not seen to co-aggregate, or only appeared to do so in a very small minority of cells (<5%).

### No obvious effect of CRMP1 on the extent of DISC1 aggregation

A previous report demonstrated that CRMP1 co-aggregates with huntingtin (HTT) in Huntington’s disease, but in doing so reduces the extent of aggregation and neurotoxicity of huntingtin

(27). It is therefore plausible that it has a similar effect on DISC1. We therefore performed a quantitative assay, in which Flag-tagged DISC1 was co-transfected into SH-SY5Y cells with EGFP alone, EGFP-CRMP1 Sv or EGFP-CRMP1 Lv, by a researcher who was blinded as to which plasmid was in each set of cells. This researcher then assessed aggregation, before having the data decoded. Co-expression with EGFP-CRMP1 Sv or Lv had no effect on the number of DISC1 aggregates per cell compared to co-expression with EGFP alone (figure 5A), although the average size of DISC1 aggregates co-expressed with CRMP1 Sv was slightly smaller (figure 5B). There was no significant difference in the number or size of co-aggregates of DISC1 with CRMP1 Sv or Lv (figure S7A-C), although levels of co-aggregates of each were higher than was seen with EGFP alone, nor was there a significant difference in number of aggregates of CRMP1 Sv compared to Lv (figure S7D). CRMP1 therefore seems to have a minimal effect on DISC1 aggregation, at least in this assay.

**Figure 5.**
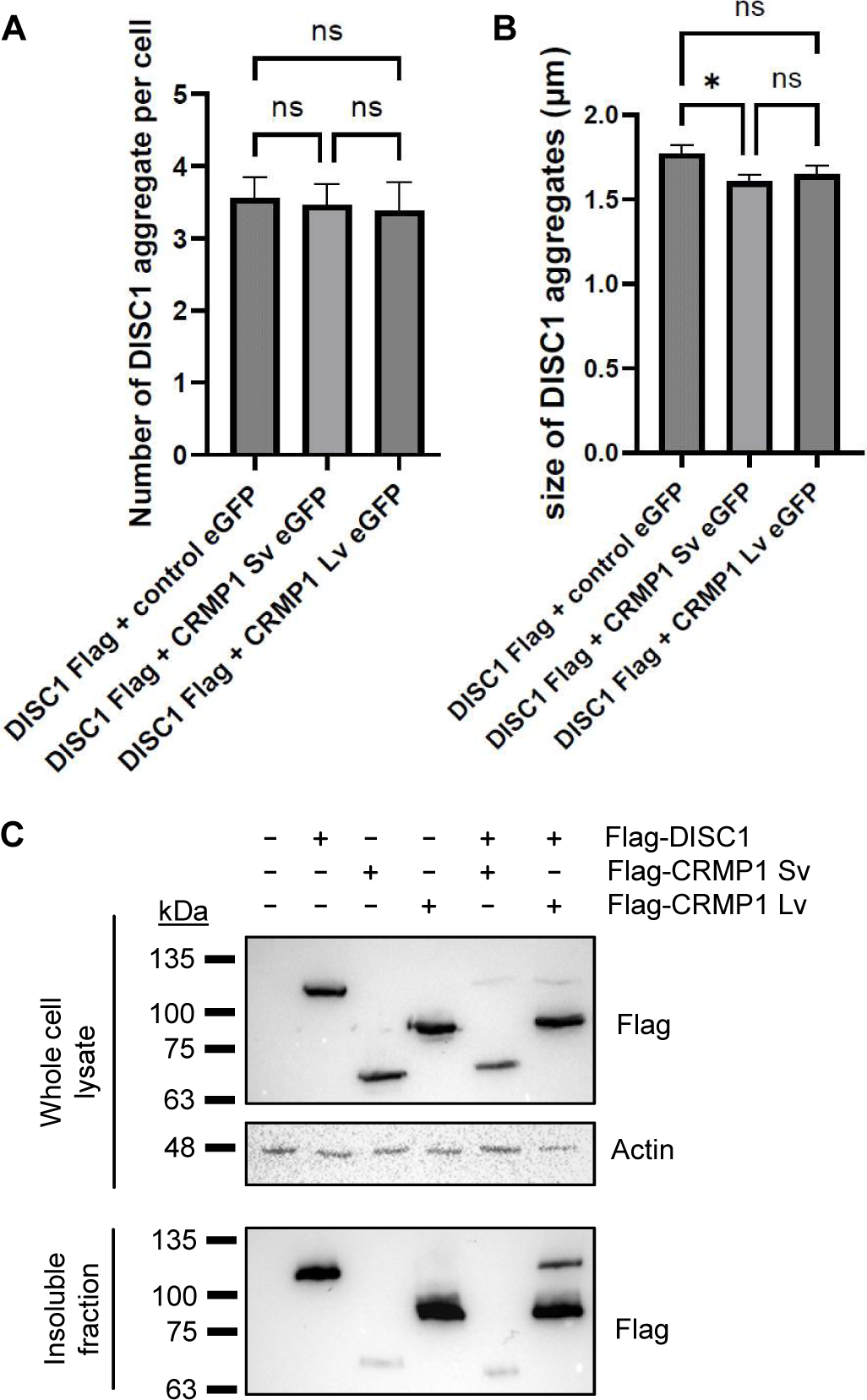
Investigating the co-aggregation of DISC1 and CRMP1. **(A-B)** Co-expression of Flag-DISC1 with EGFP, EGFP-CRMP1 Sv or EGFP-CRMP1 Lv in SH-SY5Y cells, analyzed by immunofluorescent microscopy in a blinded, quantified manner. For the purpose of this assay, an aggregate was defined as a punctate structure at least 1 µm in diameter. **(A)** Mean number of Flag (DISC1) aggregates per cell. **(B)** Mean size of DISC1 aggregates in cells displaying DISC1 aggregates. *: p < 0.05, ns: not significant, according to one-way ANOVA. **(C)** Western blots from an insoluble fraction purification assay. HEK293 cells were transfected with various combinations of Flag-tagged DISC1, CRMP1 Sv and Lv, lysed, and then had their insoluble protein fraction purified. Samples are shown from both the unfractionated cell lysate and the purified insoluble fraction, and are representative of three independent experiments.

As an alternative approach, DISC1, CRMP1 Sv and/or CRMP1 Lv were also transfected into HEK293 cells, which were subsequently lysed (all constructs were confirmed to show the same expression pattern in HEK293 as in SH-SY5Y, figure S8). These cell lysates then had their insoluble protein fraction purified, in a method directly analogous to that used on the brain samples. DISC1 and CRMP1 Lv were both prominent in the insoluble protein fraction, while CRMP1 Sv was present at a much lower level, indicating that CRMP1 Sv may have a lower aggregation propensity than DISC1 or CRMP1 Lv (figure 5C). There was no clear effect of CRMP1 on levels of insoluble DISC1.

### DISC1 can induce aggregation of TRIOBP-1, with no individual domain of TRIOBP-1 sufficient for this

To further investigate the co-aggregation of DISC1 and TRIOBP-1, we utilized our recently developed TRIOBP-1 mutant, Δ1-59Δ333-340, which lacks its optionally translated N-terminus and a short loop region in the middle of the protein (figure 6A, S9A), abolishing its ability to aggregate (6). We therefore expressed both this truncated and full length TRIOBP-1, labelled with Flag tags, together with full length DISC1 fused to EGFP. Notably, while the TRIOBP-1 mutant did not aggregate when expressed alone or with EGFP (supp. Figure S9B), it did co-aggregate with full length DISC1 in some cells (figure 6B), indicating that DISC1 can induce aggregation of TRIOBP-1. In other cells, DISC1 and the TRIOBP-1 mutant still colocalized, but without signs of aggregation (figure 6C).

**Figure 6.**
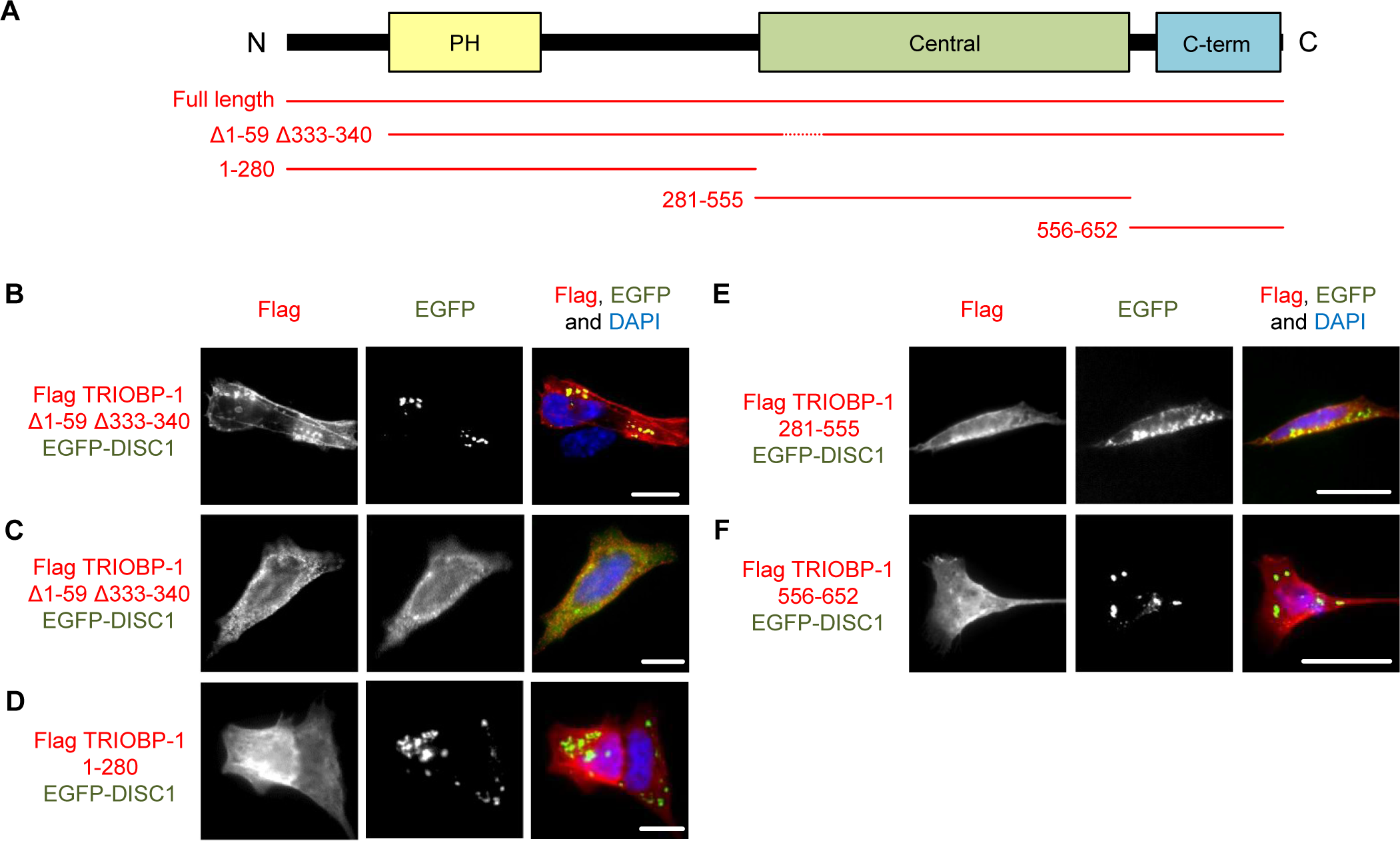
Investigating the co-aggregation of DISC1 and TRIOBP-1 in SH-SY5Y cells. **(A)** Schematic of the major domains of TRIOBP-1 (Pleckstrin homology domain, central coiled-coiled domain and C-terminal coiled-coil domain) and locations included in the plasmid vectors used here. **(B)** TRIOBP-1 Δ1-59 Δ333-340, which does not aggregate by itself, co-aggregates with DISC1 in some cells. **(C)** The same constructs in another cell, where no aggregation is seen. **(D)** The N-terminal sections of TRIOBP-1, including the PH domain, do not co-aggregate with DISC1. **(E)** The central coiled coil-domain of TRIOBP-1 shows only limited co-aggregation with DISC1. **(F)** The C-terminal coiled coil-domain of TRIOBP-1 does not co-aggregation with DISC1. All cell photos are typical of 3 or more independent experiments. Scale bars represent 10 μm.

To investigate the co-aggregation further, DISC1 was co-transfected with fragments of the TRIOBP-1 representing its disordered N-terminus and PH domain, central coiled-coil domain and C-terminal coiled-coil domain. As reported previously (19), of these only the central coiled-coil domain formed aggregates when expressed alone (figure S9A,C-E). When co-expressed with DISC1, neither the N-terminal region nor C-terminal coiled-coil domain showed signs of co-aggregation, while the central coiled-coil domain showed only weak indications compared to the full-length protein, either alone or when expressed together (figures 6D-G). It therefore appears that DISC1 needs to interact with multiple sections of the TRIOBP-1 protein in order to induce co-aggregation.

## Discussion

Impaired proteostasis, and the accumulation of insoluble protein, is an established and well recognized feature of the neurodegenerative diseases. Some diseases are characterized by a single aggregating protein, such as huntingtin in Huntington’s disease, while others are associated with a variety of distinct proteinopathies, such as in amyloid lateral sclerosis or frontotemporal lobe dementia. The reasons that these proteins begin to aggregate vary, but include both inherited and spontaneous mutations in the genes encoding them, as well as environmental stresses. Recently, various proteins have been implicated as forming aggregates in major mental illnesses, although the relationship between these proteins remains unclear.

One such protein, DISC1 has been found to aggregate in the same brain samples as two other proteins implicated in mental illness, CRMP1 and dysbindin-1 (7, 9), as well as two proteins related to neurodegenerative disease, huntingtin and TDP-43 (31, 32). In all cases, DISC1 was found to co-aggregate directly with this other protein in cell culture or other in vitro assays. It therefore could reasonably be predicted that the co-occurrence of aggregation (insoluble) proteins in the brains of mental illness proteins would typically be the results of direct co-aggregation of the proteins. In our data, however, while instances of multiple proteins aggregating in a single brain sample were relatively common, they are not high enough to imply that one protein frequently affects the aggregation state of another. Partially consistent with this, of the four proteins we investigated here, CRMP1, DISC1, NPAS3 and TRIOBP-1, only two pairs show direct co-aggregation. One of these is the previously reported CRMP1-DISC1 aggregation, the other is a novel pair: DISC1 and TRIOBP-1. We did not study dysbindin-1 here, as we were unable to get consistent antibody staining against it in our brain homogenate samples.

Co-aggregation was studied by pairwise expression of proteins in SH-SY5Y cells, one with a Flag tag and one with an EGFP fusion protein for detection. In most instances, changing which protein was in each vector did not have an effect on the results, although it was notable that CRMP1 appeared to show increased aggregation when fused to EGFP. This is potentially because the relatively large size (27 kDa) of the fusion protein increases the stability of the protein, meaning that if the protein began to misfold, it would be more likely to survive as an aggregating protein, than be destroyed by the proteasome. As a precaution, however, all pairs of proteins were expressed with both combinations of protein and plasmid vector, to minimize any effect of the EGFP fusion protein.

The ability of DISC1 and CRMP1 sv to co-aggregate was reported previously, with the two being insoluble in the same brain samples and also aggregating in cell lines (7). In this study, aggregation of CRMP1 sv was only seen when co-expressed with DISC1, and appeared more prominent than in our experiments (7), however this may be because those authors used fluorescent tags on both proteins, which can lead to heightened aggregation of CRMP1, as seen here. This study also saw GFP-tagged CRMP1 lv to aggregate (7), which matches our results, although our data showed it not to consistently aggregate without this fluorescent fusion protein. Co-aggregation of CRMP1 lv and DISC1 had not been studied before, to our knowledge, nor has the co-aggregation of the two CRMP1 isoforms, with our insolubility assays suggesting that CRMP1 Lv may be more prone to insolubility than CRMP1 Sv. This was not seen in the quantitative immunofluorescence assay, which instead looks at aggregate distribution, rather than the quantity of aggregating protein molecules.

TRIOBP-1 was first implicated as aggregating in schizophrenia using a variation of the same process used to detect insoluble CRMP1 (10), and was later validated in a distinct set of samples from schizophrenia and MDD patients (6). Aggregation of TRIOBP-1 was therefore identified separately from that of DISC1, and no interaction between the two proteins has been previously reported, to the best of our knowledge. They both however share a few common features, including expression in the brain, roles in the actin cytoskeleton (34-36) and at least two mutual protein interaction partners in NDEL1 (37-41) and TRIO (34, 42). While DISC1 could co-aggregate with full length TRIOBP-1, it could not clearly do so with individually expressed domains of TRIOBP-1, indicating that multiple points of contact between the two proteins are likely required for co-aggregation to occur. It is notable, however, that DISC1 could also co-aggregate with a mutant form of TRIOBP-1 that lacked its two aggregation-critical regions, indicating that co-aggregation of the two proteins can be driven by DISC1, and that this is a distinct mechanism from that by which TRIOBP-1 aggregates alone.

Based on this study, and consistent with previous data, it seems that these proteins implicated in mental illness can often aggregate in the brain of the same individual, however in the majority of cases this is because each protein aggregates independently (parallel aggregation). Having a direct effect of one aggregation protein on another (co-aggregation) is seemingly much rarer, and limited to certain proteins. Notably DISC1 appears to be proactive in inducing aggregation of other proteins, including TRIOBP-1, while CRMP1 co-aggregation may have a protective effect.

## Acknowledgements

This work was supported by the Croatian Science Foundation under project numbers IP-2018-01-9424, DOK-2018-09-5395 and DOK-2020-01-8580. Additional funding came from the Alexander von Humboldt Foundation through research group linkage program 1142747-HRV-IP and an equipment subsidy. The work of Éva Renner and Miklós Palkovits was supported by the Hungarian National Research, Development and Innovation Office NKFIH 2017-1.2.1-NKP-2017-00002 (National Brain Research Program NAP 2.0) and NAP2022-I-4/2022 (National Brain Research Program NAP 3.0). We thank Carsten Korth for sharing plasmids, Elizabeth Bradshaw for proof reading and the staff of the Human Brain Tissue Bank and Laboratory for their technical assistance.

