## Supplementary Material for "Co-aggregation and parallel aggregation of specific proteins in major mental illness"

**Supplementary information**

| | |
|---|---|
| Table S1: | Sources and generation of plasmid vectors used in this study |
| Table S2: | Primers used for cloning in this study |
| Figure S1: | Western blots of insoluble protein fractions, purified from human insular cortex samples |
| Figure S2: | Western blots of non-purified insular cortex sample homogenates |
| Figure S3: | Pairwise comparisons of insoluble protein levels in the insular cortex samples |
| Figure S4: | Expression patterns of Flag-tagged proteins expressed with EGFP alone and of EGFP-fused proteins |
| Figure S5: | Versions of experiments in figure 3H-K using reciprocal plasmid vectors |
| Figure S6: | Versions of experiments in figure 4A-G using reciprocal plasmid vectors |
| Figure S7: | Quantified data on CRMP1 and DISC1 co-aggregation |
| Figure S8: | Expression patterns of proteins in HEK293 match those in SH-SY5Y |
| Figure S9: | Expression patterns of TRIOBP-1 constructs, alone and with EGFP |

| No. | Vector backbone | Gene insert | Origin |
|---|---|---|---|
| 1 | pENTR1A no ccDB | (None) | Campeau et al (2009) PLOS One 4:e6529 (Addgene, clone 17398) |
| 2 | pdcDNA-FlagMyc | (None) | BCCM/LMBP Plasmid Collection, clone LMBP 4705 |
| 3 | pDEST-CMV-EGFP | (None) | Agrotis et al (2019) Autophagy 15:976-997 (Addgene, clone 123215) |
| 4 | pcDNA3.1 | CRMP1 Sv | Bader et al. (2012) Hum. Mol. Genet. 21:4406-4418 (gift from Carsten Korth) |
| 5 | pcDNA3.1 | CRMP1 Lv | Bader et al. (2012) Hum. Mol. Genet. 21:4406-4418 (gift from Carsten Korth) |
| 6 | pENTR1A | CRMP1 Sv | Subcloned from plasmid 4 in two rounds of PCR, with primers A & C, then D & E. Restriction digested and ligated into the *KpnI* and *XbaI* sites of plasmid 1. |
| 7 | pENTR1A | CRMP1 Lv | Subcloned from plasmid 5 in two rounds of PCR, with primers B & C, then D & E. Restriction digested and ligated into the *KpnI* and *XbaI* sites of plasmid 1. |
| 8 | pENTR223 | DISC1 | DNASU Plasmid Repository, clone HsCD00516321 |
| 9 | pENTR223.1 | NPAS3 | DNASU Plasmid Repository, clone HsCD00080332 |
| 10 | pENTR1A | TRIOBP-1 | Bradshaw et al (2017) J. Biol. Chem. 292:9583-9598 (gift from Carsten Korth) |
| 11 | pdcDNA-Flag | CRMP1 Sv | LR clonase recombination of plasmids 2 and 6 |
| 12 | pdcDNA-Flag | CRMP1 Lv | LR clonase recombination of plasmids 2 and 7 |
| 13 | pdcDNA-Flag | DISC1 | LR clonase recombination of plasmids 2 and 8 |
| 14 | pdcDNA-Flag | NPAS3 | Samardžija et al (2021) J. Pers. Med. 11:1070 |
| 15 | pdcDNA-Flag | TRIOBP-1 | Bradshaw et al (2017) J. Biol. Chem. 292:9583-9598 (gift from Carsten Korth) |
| 16 | pdcDNA-Flag | TRIOBP-1 Δ1-59Δ333-340 | Zaharija et al (2022) Int. J. Mol. Sci. 23:11048 |
| 17 | pdcDNA-Flag | TRIOBP-1 1-280 | Bradshaw et al (2017) J. Biol. Chem. 292:9583-9598 (gift from Carsten Korth) |
| 18 | pdcDNA-Flag | TRIOBP-1 281-555 | Bradshaw et al (2017) J. Biol. Chem. 292:9583-9598 (gift from Carsten Korth) |
| 19 | pdcDNA-Flag | TRIOBP-1 281-652 | Bradshaw et al (2017) J. Biol. Chem. 292:9583-9598 (gift from Carsten Korth) |
| 20 | pdcDNA-Flag | TRIOBP-1 556-652 | Bradshaw et al (2017) J. Biol. Chem. 292:9583-9598 (gift from Carsten Korth) |
| 21 | pDEST-CMV-N-EGFP | (Empty control) | LR clonase recombination of plasmids 1 and 4 |
| 22 | pDEST-CMV-N-EGFP | CRMP1 Sv | LR clonase recombination of plasmids 4 and 6 |
| 23 | pDEST-CMV-N-EGFP | CRMP1 Lv | LR clonase recombination of plasmids 4 and 7 |
| 24 | pDEST-CMV-N-EGFP | DISC1 | LR clonase recombination of plasmids 4 and 8 |
| 25 | pDEST-CMV-N-EGFP | NPAS3 | LR clonase recombination of plasmids 4 and 9 |
| 26 | pDEST-CMV-N-EGFP | TRIOBP-1 | LR clonase recombination of plasmids 4 and 10 |

**Table S1**. Sources and generation of plasmid vectors used in this study. See table S2 for references to primers.

| Label | Name | Primer |
|---|---|---|
| A | CRMP1-1sv-salF | GAAGTCGACATGTCGTACCAG |
| B | CRMP1-1lv-salF | GTAAGTCGACATGGCGGACCG |
| C | CRMP1-572sv-ecoR | GTAGAATTCTCATCAACCGAGGCTG |
| D | Extension F | GCTATAAGGATCCGGTACCTAGTCGACATG |
| E | Extension R | GGCACCAGCTCGAGTCTAGAATTCTCATC |

**Table S2**. Primers used for cloning in this study. “Label” corresponds to how they are referred to in table S1.

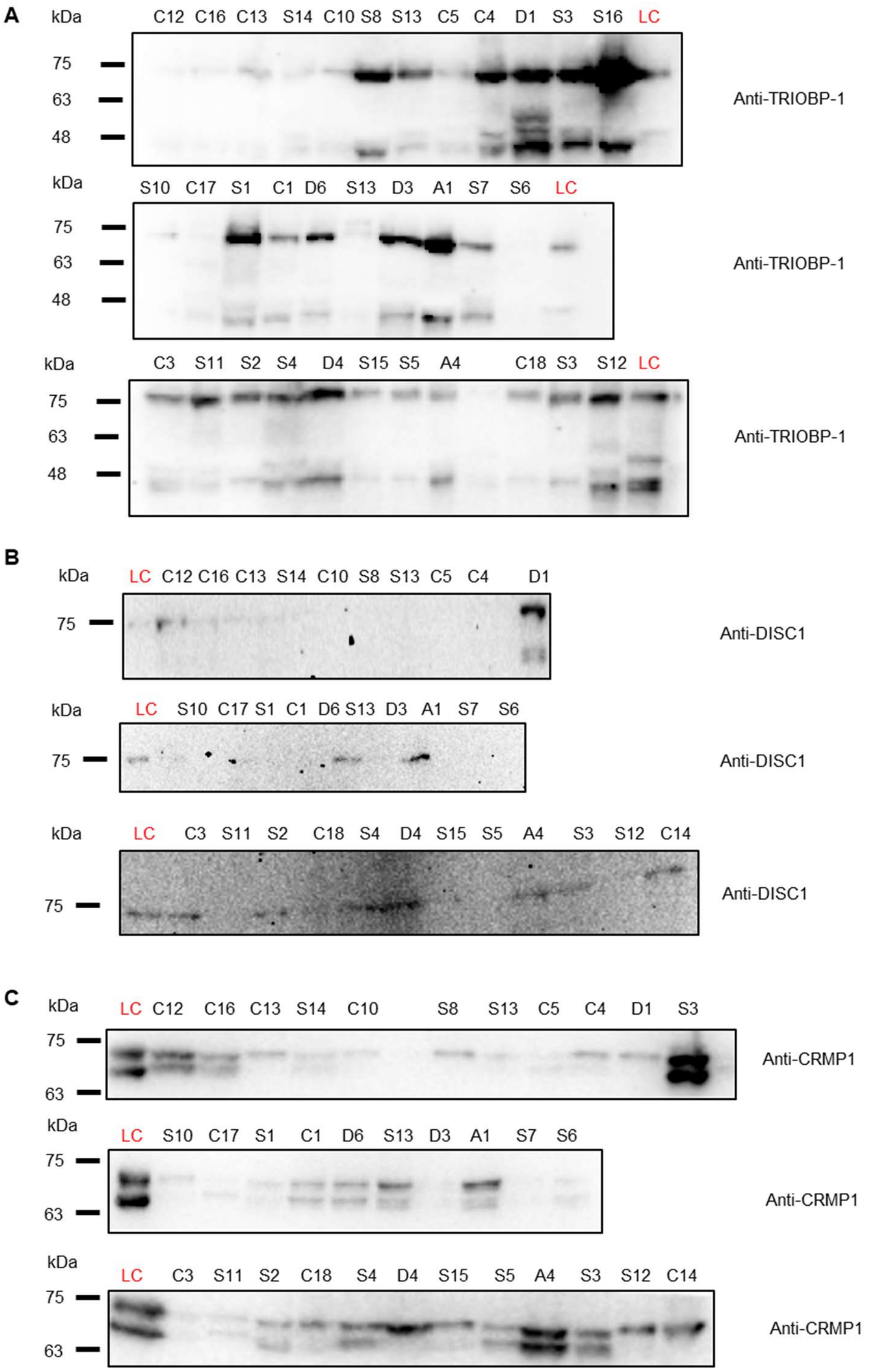


**Figure S1**. Western blots of the insoluble protein fractions, purified from insular cortex samples. Samples are labelled with diagnostic status (S: victim of suicide, C: control individual, D: major depressive disorder patient, A: Alzheimer's disease patient) and a unique number to allow comparison between blots in figures S1 and S2, which displays the corresponding non-purified homogenates. LC is a loading control: a standard sample loaded on all membranes to allow normalisation of signal quantification between gels.

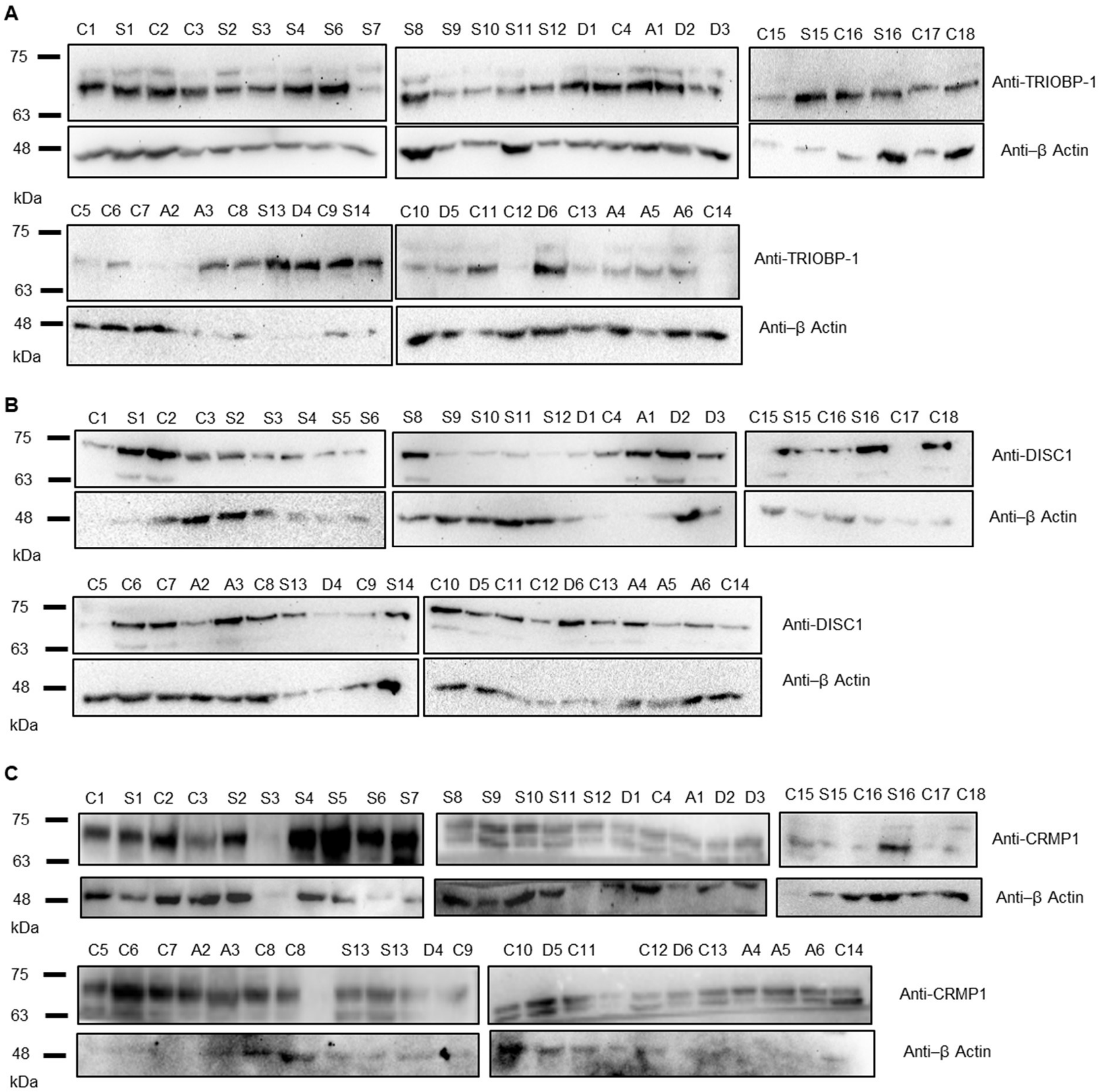


**Figure S2**. Western blots of insular cortex sample homogenates. Samples are labelled with diagnostic status (S: victim of suicide, C: control individual, D: major depressive disorder patient, A: Alzheimer's disease patient) and a unique number to allow comparison between blots in figures S1 and S2, which displays insoluble protein fractions, purified from these homogenates.

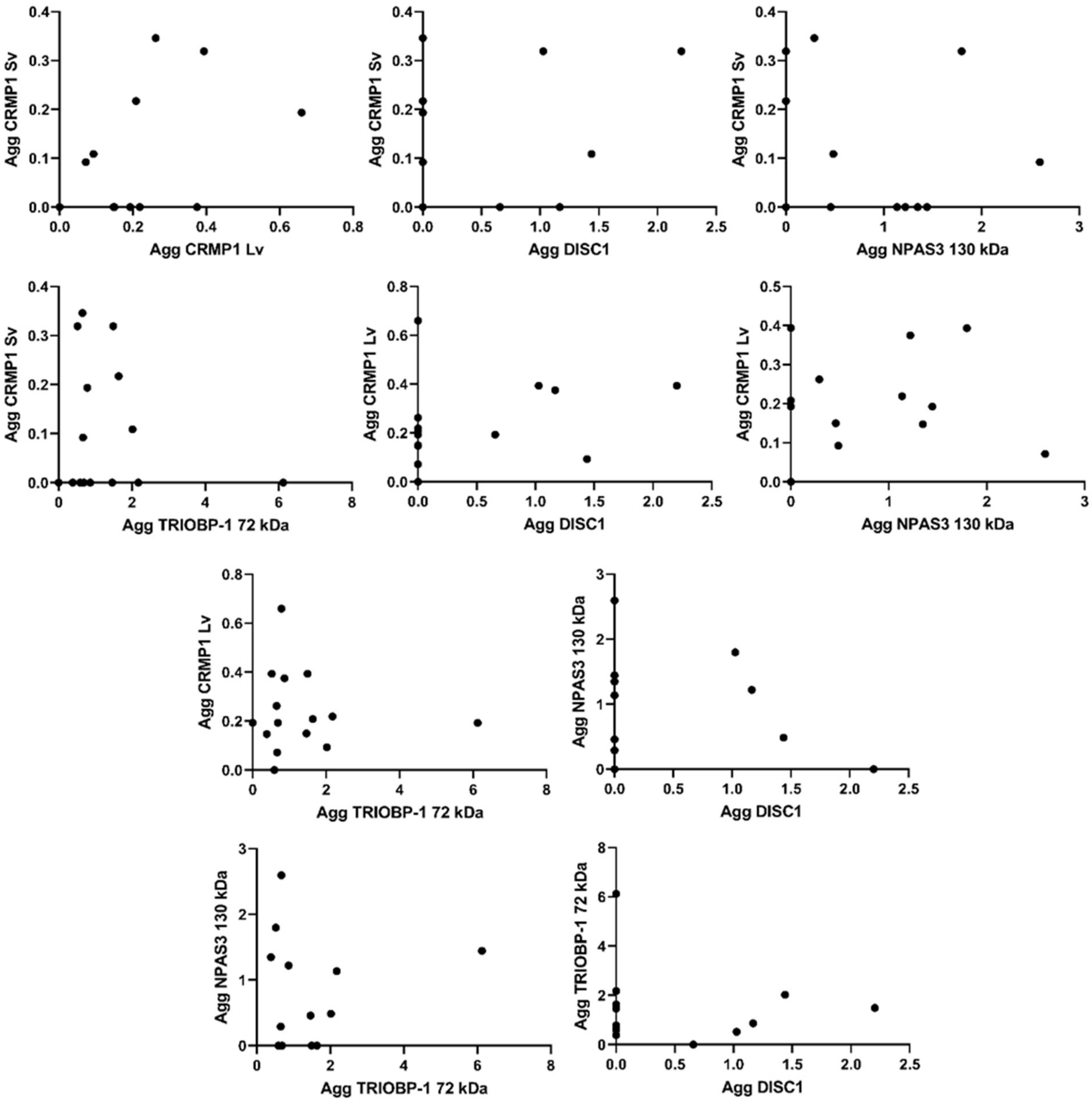


**Figure S3**. Comparison of levels of CRMP1 sv, CRMP1 lv, DISC1 (75kDa), NPAS3 (130kDa) and TRIOBP-1 in individuals. Based on data shown in figures 1 and S1.

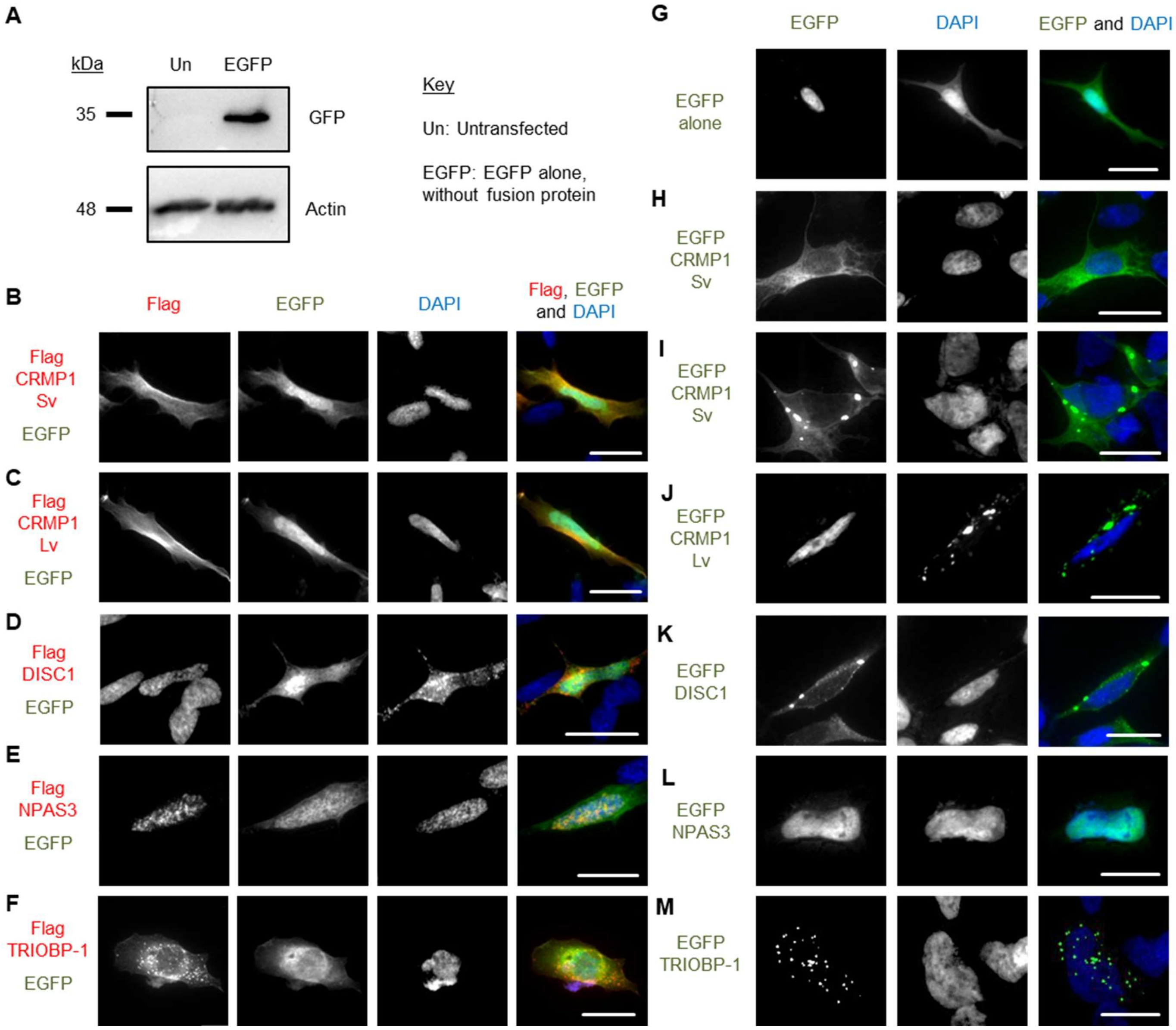


**Figure S4**. Expression of plasmids, in SH-SY5Y cells unless otherwise stated. **(A)** Western blot, showing expression of a construct encoding EGFP alone, with no fusion protein, in HEK293 cells. **(B-F)** Expression of Flag-tagged proteins (the same vectors as in figure 3C-G of the main text) together with EGFP alone: CRMP1 Sv (B), CRMP1 Lv (C), DISC1 (D), NPAS3 (E) and TRIOBP-1 (F). There are no obvious effects on expression patterns of the Flag tagged proteins. **(G)** Expression of EGFP alone. **(H-M)** Expression of EGFP-fused proteins, CRMP1 Sv with no aggregation (H, as seen in most cells), CRMP1 Sv with aggregation (I, minority of cells), CRMP1 Lv (J), DISC1 (K), NPAS3 (L) and TRIOBP-1 (M). Except for some EGFP-CRMP1 Sv & Lv cells showing aggregation (I, J), all constructs showed similar expression patterns to their Flag-tagged counterparts. All images are typical of three independent experiments. Scale bars represent 10 µm.

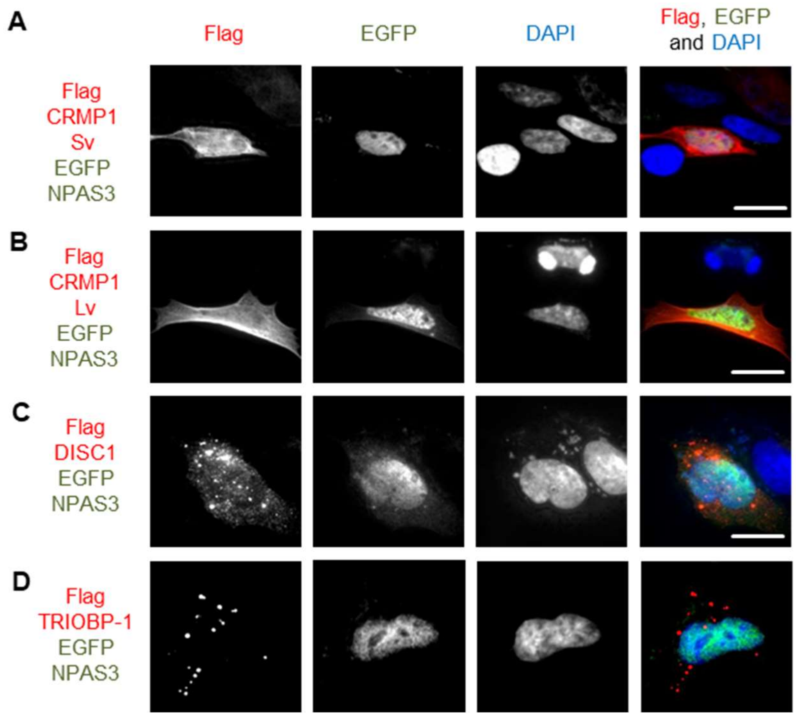


**Figure S5**. Expression of plasmids in SH-SY5Y cells, in experiments using reciprocal plasmid vectors from those in the main text. **(A)** Flag-tagged CRMP1 Sv and EGFP-fused NPAS3, neither aggregating, reciprocal experiment to that in figure 3H. **(B)** Flag-tagged CRMP1 Lv and EGFP-fused NPAS3, neither aggregating, reciprocal experiment to that in figure 3I. **(C)** Flag-tagged DISC1 and EGFP-fused NPAS3, only DISC1 is aggregating, reciprocal experiment to that in figure 3J. **(D)** Flag-tagged TRIOBP-1 and EGFP-fused NPAS3, only TRIOBP-1 is aggregating, reciprocal experiment to that in figure 3K. All images are typical of three independent experiments. Scale bars represent 10 µm.

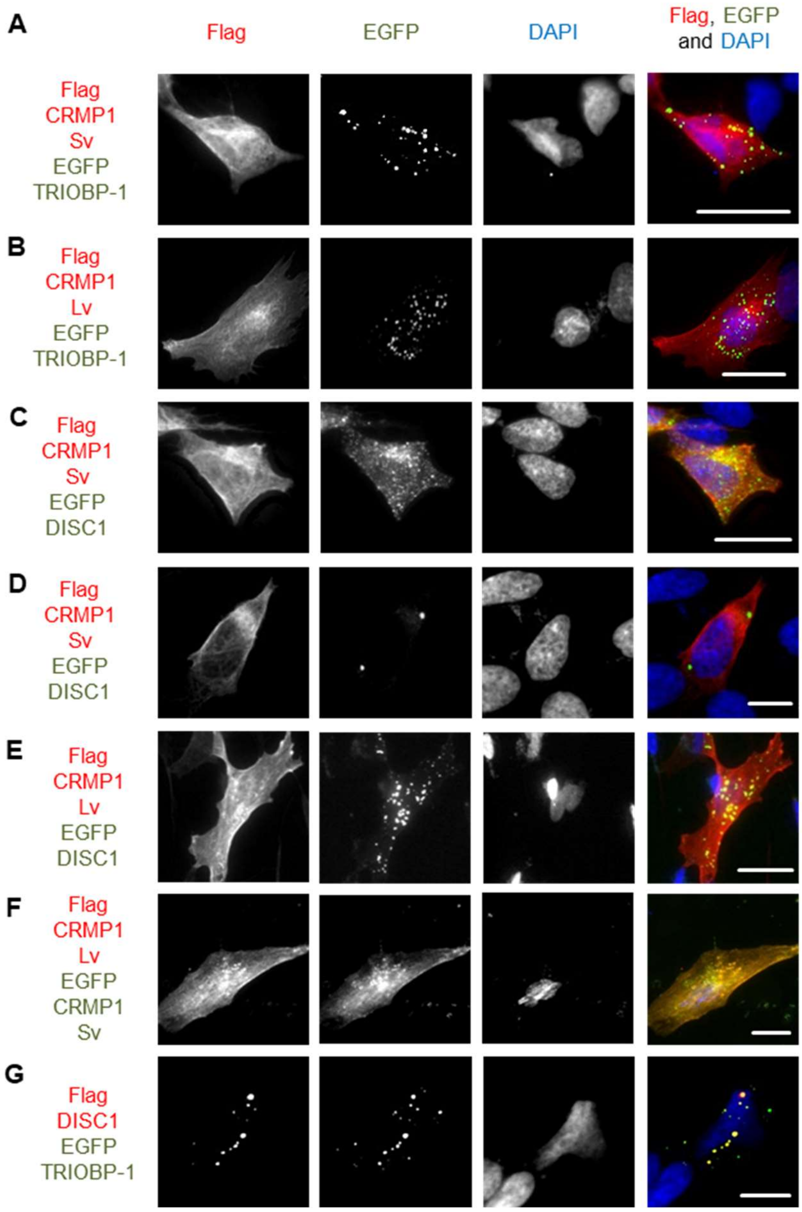


**Figure S6**. Expression of plasmids in SH-SY5Y cells, in experiments using reciprocal plasmid vectors from those in the main text. **(A)** Flag-tagged CRMP-1 Sv and EGFP-fused TRIOBP-1, only TRIOBP-1 aggregates, reciprocal experiment to that in figure 4A. **(B)** Flag-tagged CRMP-1 Lv and EGFP-fused TRIOBP-1, only TRIOBP-1 aggregates, reciprocal experiment to that in figure 4B. **(C-D)** Flag-tagged CRMP-1 Sv and EGFP-fused DISC1, example of cell with co-aggregation (C) and DISC1 aggregation only (D), reciprocal experiment to that in figure 4C-D. **(E)** Flag-tagged CRMP-1 Lv and EGFP-fused DISC1 with some co-aggregation, reciprocal experiment to that in figure 4E. **(F)** Flag-tagged CRMP1 Lv and EGFP-fused CRMP1 Sv, with co-aggregation seen, reciprocal experiment to that in figure 4F. **(G)** Flag-tagged DISC1 and EGFP-fused TRIOBP-1, with extensive co-aggregation seen, reciprocal experiment to that in figure 4G,H. All images are typical of three independent experiments. Scale bars represent 10 µm.

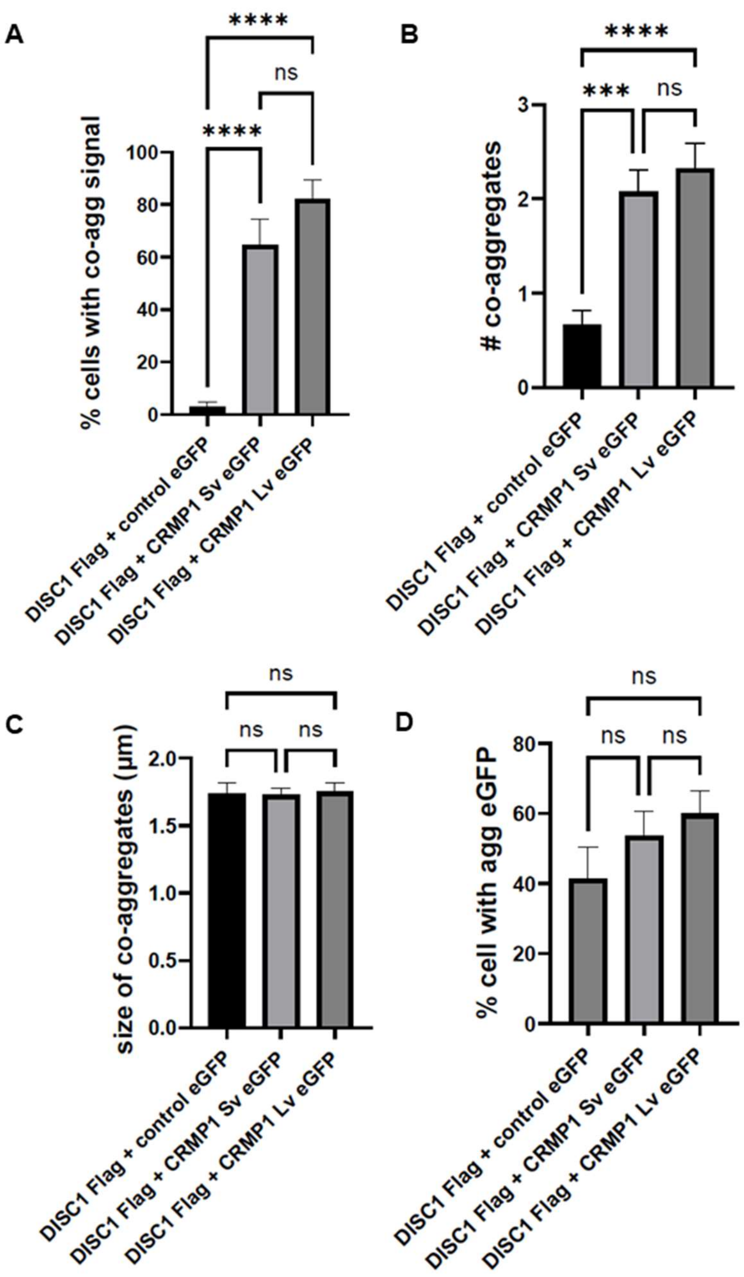


**Figure S7**. Additional quantification of DISC1 and CRMP1 aggregation in a co-aggregation blinded immunofluorescent microscopy assay (see figure 5). SH-SY5Y cells were transfected with Flag-DISC1 and one of EGFP, EGFP-CRMP1 Sv or EGFP-CRMP1 Lv. **(A)** Mean percentage of successfully transfected cells in which co-aggregation of Flag and EGFP seem to occur, defined as co-localisation at a punctate structure at least 1 µm in diameter. **(B)** Number of distinct co-aggregates in such cells. **(C)** Mean size of co-aggregates in such cells. **(D)** Percentage of successfully transfected cells in which aggregates of EGFP were seen (regardless of DISC1 aggregation status). All statistics are one-way ANOVA: ****:  $p < 0.0001$, ***: $p < 0.001$, ns: not significant.

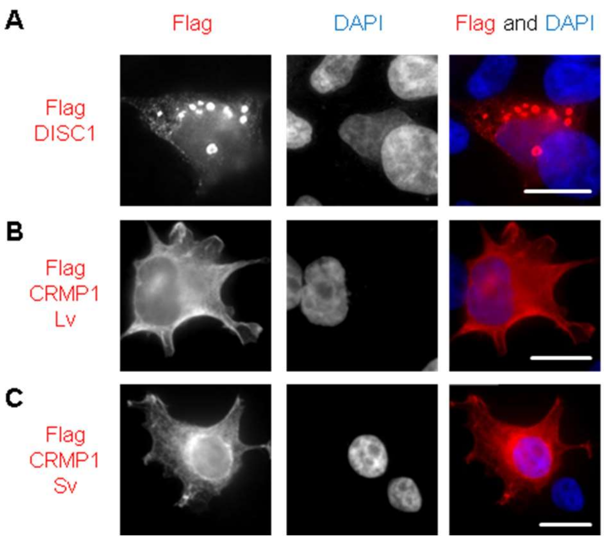


**Figure S8**. Expression patterns of Flag-tagged proteins expressed in HEK293 closely matches the patterns seen when the same proteins are expressed in SH-SY5Y (as shown in figure 3). **(A)** DISC1. **(B)** CRMP-1 Lv. **(C)** CRMP-1 Sv. Scale bars represent 10 µm.

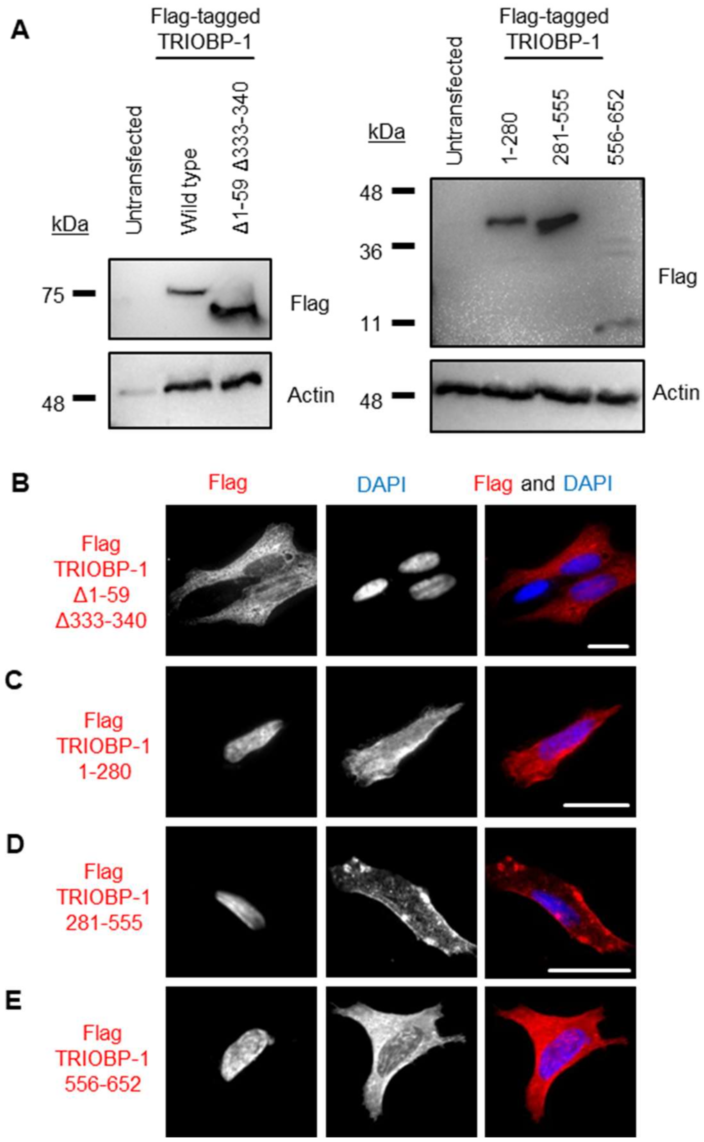


**Figure S9**. Expression patterns of Flag-tagged TRIOBP-1 constructs, in SH-SY5Y cells unless otherwise stated. **(A)** Western blots of HEK293 cells expressing the various TRIOBP-1 fragment. **(B-E)** Expression of Flag-tagged TRIOBP-1 proteins, expressed alone in cells: (B) Non-aggregating TRIOBP-1 lacking amino acids 1-59 and 333-340, (C) Amino acids 1-280, encoding the PH domain and unstructured/disordered regions near the N-terminus, (D) amino acids 281-555, encoding the central coiled-coil domain, (E) amino acids 555-652, encoding the C-terminal coiled-coil domain.
